# Spotty distributions: Spotted Gar (*Lepisosteus oculatus*) and Spotted Sucker (*Minytrema melanops*) range expansion in eastern Lake Erie

**DOI:** 10.1101/2023.02.08.527752

**Authors:** Daniel J. MacGuigan, Isabel Porto-Hannes, Brian Foote, Nathan J. C. Backenstose, Christopher Osborne, Kimberly Louisor, Hannah Waterman, Sarah L. Chang, Jacob L. Cochran, Trevor J. Krabbenhoft

## Abstract

Natural range expansions in warm-water freshwater fishes are currently not well understood, but shifts in native species distributions can be influenced by many factors, including habitat restoration or degradation and climate change. Here, we provide empirical evidence of range expansions observed in two native freshwater fish species in Lake Erie: the Spotted Gar (*Lepisosteus oculatus)* and Spotted Sucker (*Minytrema melanops*). We confirmed our field identifications of *L. oculatus* and *M. melanops* using mtDNA barcoding. Maximum likelihood phylogenetic analyses reveal that our samples confidently resolve in the *L. oculatus* and *M. melanops* clades respectively, with additional identification support from BLAST searches. Notably, we found no correlation between the increased detection rate of both species and an increase in sampling effort when compared to previous records. Historically, eastern Lake Erie experienced habitat degradation through channelization, siltation, dredging, and toxification of sediments. We hypothesize that recent habitat remediation efforts have provided suitable habitat for both species to recolonize shallow waters with densely vegetated habitat (>90% substrate coverage). Both species are likely to continue their northern expansion as habitats are restored and climatic changes favor warm-water fishes.

## INTRODUCTION

Distributions of freshwater fishes are shaped by a combination of physiological, ecological, and geological factors and are often reflective of the evolutionary history of a species (Jackson et al., 2001). However, these factors are rarely static, and species distributions expand and contract over time (Jackson et al., 2001). Range shifts can also be influenced by anthropogenic effects such as human-mediated climate change (Comte et al., 2013; Lynch et al., 2016b; Myers et al., 2017), habitat destruction (Scheuerell and Schindler, 2004), damming of migratory streams for management of freshwater use (Barbarossa et al., 2020; Ormerod et al., 2010), and deliberate and unintentional transport of species outside of their native distributions (Rahel, 2002). These human-induced changes can drastically alter the regional and global distributions of freshwater fishes on a contemporary time scale (Su et al., 2021; Wheeler et al., 2020). Warm-water fish species in lower latitudinal areas are also expected to shift towards higher latitudes in response to climate variation (Last et al., 2011). Due to the high ecological, economic, and cultural importance of freshwater fishes (Lynch et al., 2016a), documenting and monitoring changes in species distributions remains an important task in freshwater ecosystems experiencing climactic shifts (Ebersole et al., 2020; Olden et al., 2010).

The Laurentian Great Lakes (hereafter, Great Lakes) system is ideal for examining the impacts of climactic and other anthropogenic changes on species distributions. The Great Lakes are connected by both natural and human-constructed waterways and represent the largest system of freshwater lakes by surface area and the second largest by volume (Government of Canada and US EPA, 1995). The Great Lakes are also relatively young, having been completely glaciated as recently as 17,400 years ago (Dalton et al., 2020). Although the Great Lakes are a well-known hotbed of invasion by non-native species (DeRoy and MacIsaac, 2020; Ricciardi and MacIsaac, 2000), the connectivity and young age of these lakes might also facilitate distribution shifts for native species. In the context of invasions and changing community dynamics, Lake Erie is notable among the Great Lakes for several reasons. Lake Erie is the shallowest and warmest of the Great Lakes, and therefore perhaps the most easily perturbed by nutrient influxes (Scheffer et al., 1993; Scheffer and van Nes, 2007). Lake Erie also sits at a junction between the Mississippi River basin (which drains to the Gulf of Mexico) and river basins that drain to the Atlantic Ocean, providing a contact point between the disparate fish fauna of these drainages (April et al., 2013). Lake Erie historically had a larger commercial fishery than all the other Great Lakes combined (Regier and Hartman, 1973). It experienced extensive pollution and habitat destruction up until the 1970s (George et al., 2022b; Koonce et al., 1996; Regier and Hartman, 1973), and the Buffalo River and the upper Niagara River in the eastern Lake Erie basin are both listed as Areas of Concern (AOCs) under the Great Lakes Water Quality Agreement of 1972 (GLWQA, 1972). These factors make Lake Erie a likely candidate region for range expansions and contractions of freshwater fishes.

Two fish species that are historically absent or rare in eastern Lake Erie are the Spotted Gar (*Lepisosteus oculatus)* and the Spotted Sucker (*Minytrema melanops)*. Both of these fishes are widespread throughout the southern and central United States (Page and Burr, 2011) but are sporadically distributed in the Great Lakes basin (Boothroyd et al., 2016; Cudmore-Vokey and Crossman, 2000; George et al., 2022a; Hubbs and Lagler, 2004; Trautman, 1981). *Lepisosteus oculatus* is a large (up to 80 cm) top-level piscivorous predator preferring clear vegetated waters, particularly in wetlands and floodplain habitat of lakes and large rivers (Hubbs and Lagler, 2004; Page and Burr, 2011; Snedden et al., 1999; Suttkus, 1963; Trautman, 1981; Walker et al., 2013), while young-of-year individuals prefer low turbidity, shallow, vegetated nearshore habitat (McAllister et al., 2022). Adult *L. oculatus* have substantial dispersal capabilities, with reports of movement speeds up to 40.1 m/h (Snedden et al., 1999). In contrast, *M. melanops* is a smaller sized species (up to 60 cm) that largely feeds on benthic crustaceans and zooplankton (Hubbs and Lagler, 2004; White and Haag, 1977) and requires swift riffles with gravel or sand substrate for spawning (White, 1974; Mcswain and Gennings, 1972). Extensive siltation and channelization have led to historical declines in abundance in both *L. oculatus* (Gray et al., 2012), and *M. melanops* across the northern extent of their ranges (White, 1974; Trautman, 1981).

Despite their different life histories, *L. oculatus* and *M. melanops* have recently undergone range expansions in Lake Erie. Here we report the first records of the *L. oculatus* in the eastern basin of Lake Erie in New York, which likely represents an established population. In addition, we report a significant population expansion of the *M. melanops* in eastern Lake Erie. We hypothesize that these are natural range expansions (versus, e.g., baitbucket introductions) and were driven by a combination of human-induced climate change and restoration of natural habitats. *Lepisosteus oculatus* and *M. melanops* are likely to continue their northward range expansion and highlight the importance of continued monitoring throughout the Great Lakes basin.

## MATERIALS AND METHODS

We conducted sampling in the New York waters of Lake Erie and Niagara River between May-October 2022. Electrofishing was conducted on a boat mounted electro-fisher Infinity HC-80 Midwest Lake Electrofishing Systems following University at Buffalo IACUC Protocol 201900161. Water temperature and ambient conductivity were collected at each site to determine appropriate power (amperage expand). We sampled 500 m long transects for 20 minutes each. Fish were collected by dipnet and were photographed, measured, and pectoral fin clips were taken and stored in 95% ethanol at −20º C until DNA extraction, and fish were released at the same location where originally caught. All individuals were identified in the field using morphological characters. *Lepisosteus oculatus* key characters were large spots on the head and body and a shorter and broader jaw in contrast to Longnose Gar, *L. osseus* (Hubbs and Lagler, 2004). *Minytrema melanops* is characterized by an inferior, protrusible mouth and parallel rows of conspicuous dark spots on the dorsum and sides (Hubbs and Lagler, 2004). One *L. oculatus* individual (TJK 29) was euthanized following UB IACUC Protocol 201900161, fixed in 10% formalin, and deposited at the Yale Peabody Museum of Natural History (YPM ICH.035837).

To create range maps for *L. oculatus*, occurrence records were downloaded from FishNet2, VertNet, and GBIF. We included only records from museum collections, state agencies, or federal agencies. Occurrence records in Canada were obtained from COSEWIC (Mandrak and Bouvier, 2014). Records were overlain on a modified range map from David and Wright (2017) and Page and Burr (2011). Range maps were created using ArcGIS Pro 3.0.

We sequenced the mitochondrial gene cytochrome b (*cytb)* to confirm our field identification of *L. oculatus* and *M. melanops*. Genomic DNA was extracted using PureLink Genomic DNA Mini Kit (Invitrogen), following standard protocols. Polymerase chain reaction (PCR) was performed using LongAmp Taq 2X Master Mix (New England Biolabs) and previously described primers GLU (5’ GACTTGAAGAACCACCGTTG 3’) and THR (5’ TCCGACATTCGGTTTACAAG 3’) (Near et al., 2000). PCR reaction conditions were as follows: initial denaturation at 94 °C for 180 s, 30 cycles of denaturation at 94 °C for 30 s, annealing at 55 °C for 30 s, and extension at 72 °C for 90 s, then a final extension at 72 °C for 300 s. Amplified double-stranded DNA was purified using 1.8X AmpureXP beads and two washes with 70% ethanol. Forward and reverse amplified DNA strands were sequenced (Sanger method) by Azenta/GENEWIZ (Azenta US, Inc., South Plainfield, NJ, USA). Raw reads were trimmed by eye using chromatograms and assembled into contiguous sequences using Geneious Prime v.2019.2.3 (http://www.geneious.com).

Sequences of the mitochondrial DNA gene *cytb* were downloaded from NCBI Genbank for Lepisosteidae (n=18) and Catostomidae (n=35) (Supplementary Table 1, Supplementary Table 2). Our assembled sequences of *cytb* were combined with Genbank sequences and aligned with MAFFT (Katoh and Standley, 2013). Phylogenetic analyses were performed using W-IQ-TREE (Trifinopoulos et al., 2016). Best fit codon partition models for both data sets were determined based on Bayesian Information Criterion (BIC) scores (Kalyaanamoorthy et al., 2017). Phylogenetic tree reconstruction of *cytb* sequences for each family were completed using maximum likelihood (Minh et al., 2020) and the best fit substitution models, with 1,000 ultrafast bootstrap replicates to assess topological support (Hoang et al., 2018). All samples for each species had the same *cytb* haplotype; therefore, we selected one representative for *L. oculatus* (TJK 05) and *M. melanops* (TJK 19) to compare with the NCBI nt database using the Basic Local Alignment Search Tool (BLASTn) with default parameters. Uncorrected pairwise p-distances for the *cytb* sequences of *M. melanops* were calculated using the R package ape v.5.6-2 (Paradis et al., 2004).

## RESULTS AND DISCUSSION

### Lepisosteus oculatus range expansion in eastern Lake Erie

Four *L. oculatus* individuals were collected over a three-month period at localities ranging from Dunkirk Harbor (Lake Erie) to the Upper Niagara River (Table 1, Fig. 1, 2). The first specimen (TJK 05, Table 1, Fig. 2), was collected in the Niagara River on June 13^th^, 2022, in shallow water ranging between 0.3 – 1 m deep. The total length (TL) of this specimen measured 521.21 mm. The second specimen (TJK 20, Table 1, Fig. 2, 495.20 mm TL) was collected in approx. 2 m deep water on June 28^th^, 2022, in Lake Erie, Buffalo Harbor. The third and fourth specimens were collected from the same area in Lake Erie, Dunkirk Harbor (Table 1, Fig. 2). Specimen TJK 29 was collected July 19^th^, 2022, measuring 501.00 mm TL, while TJK 36 was caught August 11^th^, 2022, measuring 523.72 mm TL (Table 1, Fig. 2). Water depth for the Dunkirk Harbor at the collection sites ranged between 0.3 – 1 m deep. Overall, all *L. oculatus* individuals were found in heavy aquatic vegetated habitat largely consisting of eelgrass (*Vallisneria americana*) covering > 90% of the substrate in low flow areas.

**Figure 1.**
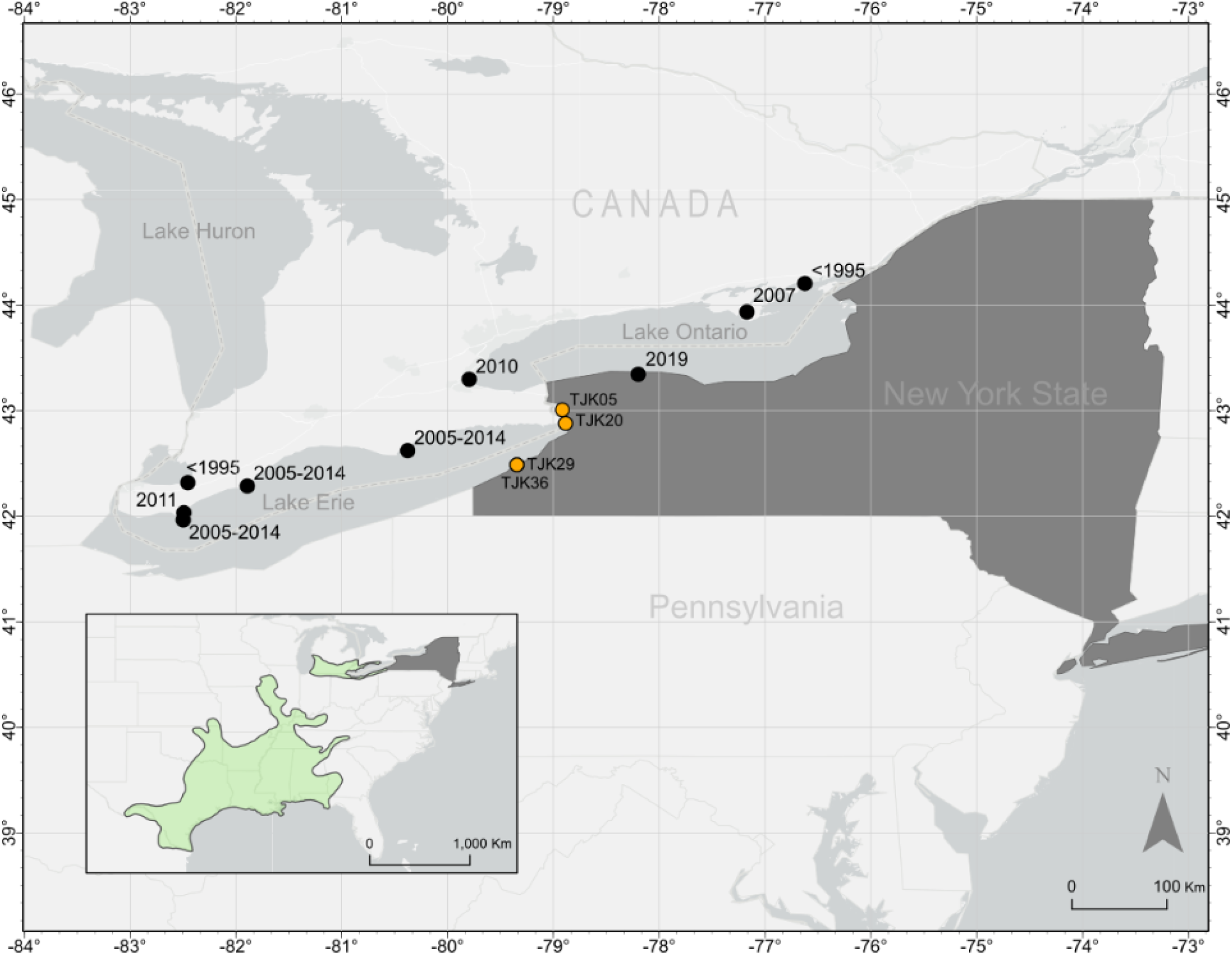
*Lepisosteus oculatus* range map. Inset map shows the native range (green) of *L. oculatus*, modified from David and Wright (2017) and Page and Burr (2011). Orange points indicate new *L. oculatus* records reported here.

**Figure 2.**
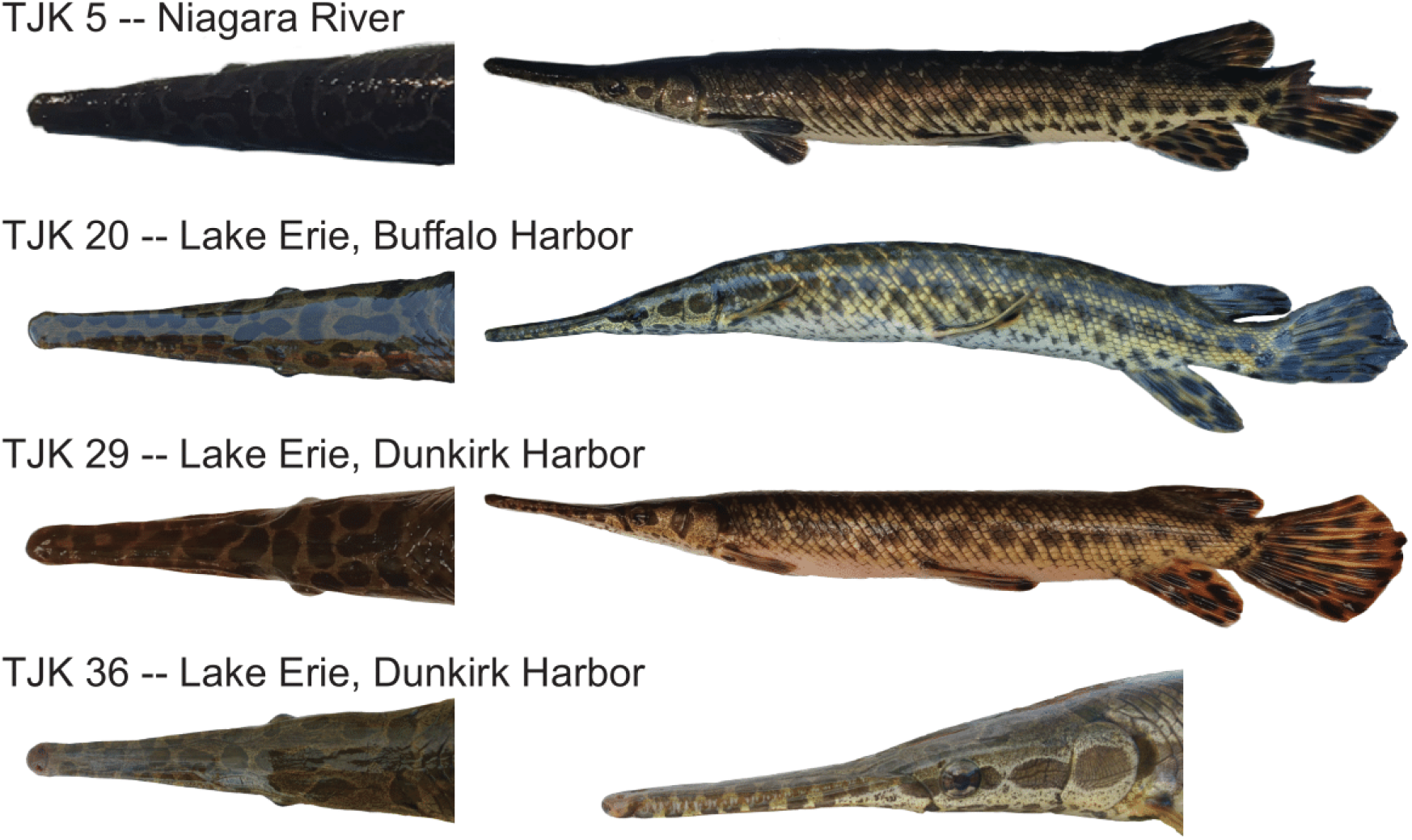
*Lepisosteus oculatus* individuals from eastern Lake Erie, with dorsal views of the head and lateral views of the left side.

**Table 1.** Sampling localities for *Lepisosteus oculatus* and *Minytrema melanops* in Lake Erie and the Niagara River, NY, USA. NA: field observations, no tissue samples were taken.

| Species | Tissue Number | Locality | Collection Date | Latitude | Longitude | Collectors |
| --- | --- | --- | --- | --- | --- | --- |
| *Lepisosteus oculatus* | TJK29 | Dunkirk Harbor | 7/19/2022 | N 42.48832 ° | W 79.34523 ° | USFWS |
|  | TJK36 | Dunkirk Harbor | 8/11/2022 | N 42.48832 ° | W 79.34523 ° | USFWS, B. Foote, I. Porto-Hannes |
|  | TJK20 | Buffalo Harbor | 6/28/2022 | N 42.88063 ° | W 78.88521 ° | C. Krabbenhoft, B. Foote, I. Porto-Hannes |
|  | TJK05 | Niagara River | 6/13/2022 | N 43.07142 ° | W 78.94587 ° | H. Waterman, B. Foote, I. Porto-Hannes |
| *Minytrema melanops* | TJK07 | Buffalo Harbor | 6/14/2022 | N 42.86005 ° | W 78.87530 ° | H. Waterman, K. Louisor, B. Foote |
|  | TJK19 | Buffalo Harbor | 6/28/2022 | N 42.87107 ° | W 78.88424 ° | C. Krabbenhoft, B. Foote, I. Porto-Hannes |
|  | TJK21 | Buffalo Harbor | 6/28/2022 | N 42.88123 ° | W 78.88418 ° | C. Krabbenhoft, B. Foote, I. Porto-Hannes |
|  | TJK22 | Buffalo Harbor | 6/28/2022 | N 42.88123 ° | W 78.88418 ° | C. Krabbenhoft, B. Foote, I. Porto-Hannes |
|  | TJK23 | Buffalo Harbor | 6/28/2022 | N 42.88265 ° | W 78.88538 ° | C. Krabbenhoft, B. Foote, I. Porto-Hannes |
|  | TJK24 | Buffalo Harbor | 6/28/2022 | N 42.88265 ° | W 78.88538 ° | C. Krabbenhoft, B. Foote, I. Porto-Hannes |
|  | NA | Buffalo Harbor | 6/21/2022 | N 42.86005 ° | W 78.87530 ° | H. Waterman, B. Foote, I. Porto-Hannes |
|  | NA | Buffalo Harbor | 6/28/2022 | N 42.87107 ° | W 78.88424 ° | C. Krabbenhoft, B. Foote, I. Porto-Hannes |
|  | NA | Buffalo Harbor | 7/7/2022 | N 42.87989 ° | W 78.88471 ° | H. Waterman, B. Foote, K. Louisor I. Porto-Hannes |
|  | NA | Dunkirk Harbor | 7/19/2022 | N 42.48905 ° | W 79.34164 ° | H. Waterman, B. Foote, I. Porto-Hannes |
|  | NA | City Ship Canal/Buffalo River | 8/15/2022 | N 42.86031 ° | W 78.86986 ° | B. Foote, K. Louisor, I. Porto-Hannes |
|  | NA | Buffalo Harbor | 8/25/2022 | N 42.86005 ° | W 78.87530 ° | B. Foote, K. Louisor, I. Porto-Hannes |
|  | NA | City Ship Canal/Buffalo River | 9/8/2022 | N 42.86031 ° | W 78.86986 ° | H. Waterman, B. Foote, I. Porto-Hannes |
|  | NA | Buffalo Harbor | 9/20/2022 | N 42.87989 ° | W 78.88471 ° | S. Chang, B. Foote, I. Porto-Hannes |
|  | NA | Buffalo Harbor | 10/6/2022 | N 42.84964 ° | W 78.86657 ° | S. Chang, L. Gray, B. Foote, I. Porto-Hannes |
|  | NA | City Ship Canal/Buffalo River | 10/6/2022 | N 42.85717 ° | W 78.86742 ° | S. Chang, L. Gray, B. Foote, I. Porto-Hannes |
|  | NA | City Ship Canal/Buffalo River | 10/6/2022 | N 42.85031 ° | W 78.86986 ° | S. Chang, L. Gray, B. Foote, I. Porto-Hannes |
|  | NA | Buffalo Harbor | 10/11/2022 | N 42.84735 ° | W 78.86488 ° | S. Chang, B. Foote, I. Porto-Hannes |

We sequenced 1,137 bp of the mitochondrial DNA gene *cytb* for the *L. oculatus* samples from the eastern Lake Erie basin. All specimens had identical haplotypes. Although most BLASTn hits for the *L. oculatus cytb* haplotype matched *L. oculatus* samples (Table 2), the top hit was an individual labeled as *L. osseus* (DQ536423.1). Our phylogenetic analyses (Fig. 3) resolved the same backbone topology for *Lepisosteus* as previous studies (Wright et al., 2012). All *Lepisosteus* species were monophyletic with two exceptions: one *L. oculatus* individual (NC_004744, Inoue et al 2003) resolves with *L. platyrinchus* and one *L. osseus* sample (NC_008104, Broughton and Reneau, 2006) resolves with *L. oculatus* (Fig. 3). Hybridization among gars is not uncommon (Herrington et al., 2008; Lyons and Sipiorski, 2020), so these misplaced individuals may represent mitochondrial capture via introgression. Alternatively (and perhaps more likely), these two individuals may have been misidentified. Unfortunately, no voucher specimens are associated with the problematic sequences. The gar samples from the eastern Lake Erie basin were confidently resolved in the *L. oculatus* clade.

**Figure 3.**
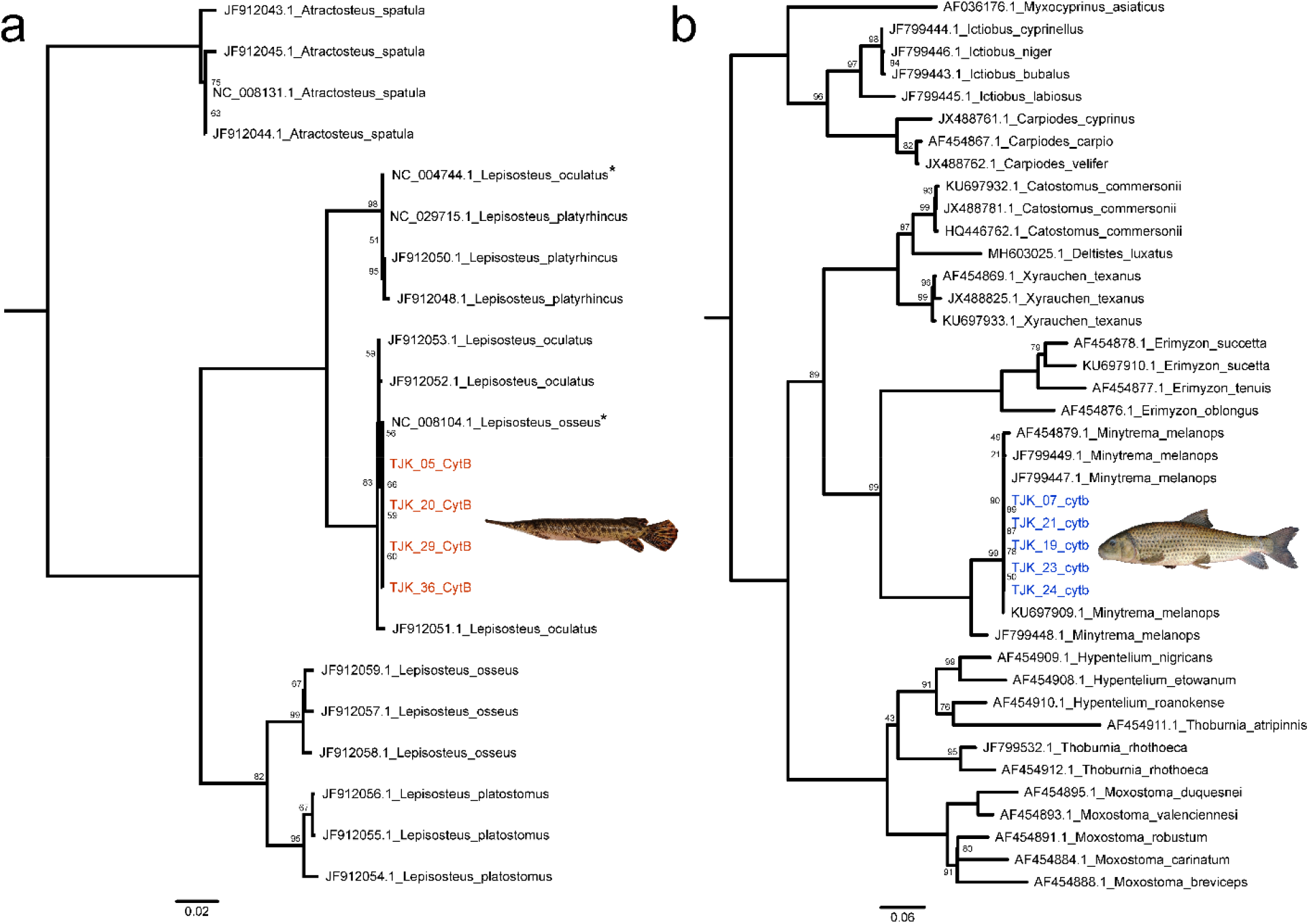
*Cytb* maximum likelihood phylogenies for a) gars and b) suckers. Branch lengths are proportional to the expected number of substitutions per site. Nodes with 100% bootstrap support are unlabeled. Individuals reported here from eastern Lake Erie are colored red (*Lepisosteus oculatus*) or blue (*Minytrema melanops*). Asterisks indicate two gar sequences downloaded from GenBank that are likely misidentified.

**Table 2.**
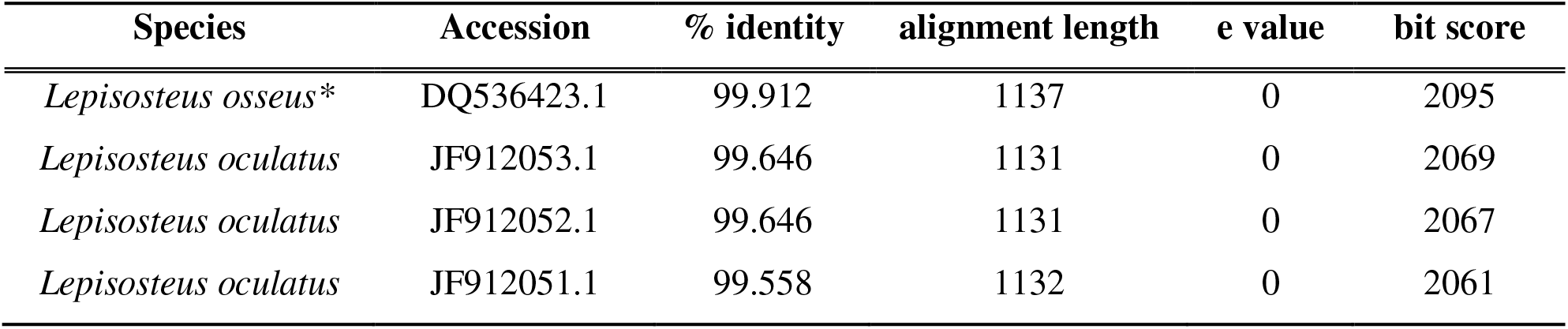
Top four BLAST hits from *Lepistosteus oculatus* sample TJK 05. The asterisk indicates a sample from GenBank that was likely misidentified (see Fig. 2).

| Species | Accession | % identity | alignment length | e value | bit score |
| --- | --- | --- | --- | --- | --- |
| *Lepisosteus osseus* * | DQ536423.1 | 99.912 | 1137 | 0 | 2095 |
| *Lepisosteus oculatus* | JF912053.1 | 99.646 | 1131 | 0 | 2069 |
| *Lepisosteus oculatus* | JF912052.1 | 99.646 | 1131 | 0 | 2067 |
| *Lepisosteus oculatus* | JF912051.1 | 99.558 | 1132 | 0 | 2061 |

The native range of *L. oculatus* is disjunct, with a northern distribution extending from southwestern Michigan to the Pennsylvania-New York border (Fig. 1). The peripheral Great Lakes basin populations are hypothesized to have split from the southern “core” populations approximately 8,000 years ago (Bailey and Smith 1981, Hocutt and Wiley 1986, Hubbs et al. 2004). The northern populations have much lower genetic diversity (David and Wright, 2017; Glass et al., 2015) and exhibit counter gradient variation, such as faster growth (David et al., 2015) compared to southern populations. The new records reported here show that peripheral populations of *L. oculatus* have extended the eastern boundary of their range into New York waters. There have been previously published sightings of *L. oculatus* further northeast in Lake Ontario (Cudmore-Vokey and Crossman, 2000), including one recent record from New York (George et al., 2022a). However, it was unclear whether these records represented stray individuals or established populations. The presence of *L. oculatus* in Lake Ontario is likely to increase as the eastern Lake Erie peripheral population expands and makes use of human-constructed waterways connecting lakes Ontario and Erie, namely the Welland and Erie canals. We hypothesize that the warmer waters and increasing vegetation provide suitable habitat for *L. oculatus* to continue its range expansion.

### Population expansion of Minytrema melanops in eastern Lake Erie

*Minytrema melanops* was observed and collected at several localities in eastern Lake Erie and the Buffalo River (Table 1) which consist of coastal urban systems. Most collection sites were characterized by shallow waters with dense submerged aquatic vegetation, similar to the habitat to where *L. oculatus* was observed (Supplementary Table 3). However, *M. melanops* was also found at greater depths (∼3.1 m) in marinas (e.g., Dunkirk Harbor, Erie Basin Marina, and Buffalo Small Boat Harbor, NY).

**Table 3.**
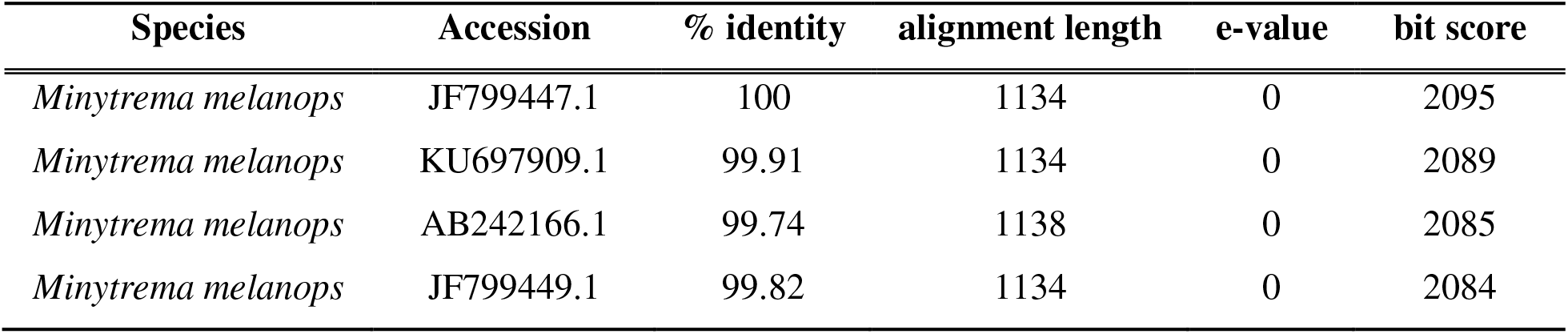
Top four BLAST hits for *Minytrema melanops* sample TJK 24.

We sequenced 1,137 bp of the mitochondrial DNA gene *cytb* for the *M. melanops* samples from the eastern Lake Erie basin. Like *L. oculatus*, all *M. melanops* specimens had identical haplotypes. BLASTn searches unambiguously confirmed the identity of our *M. melanops* samples (Table 3). Maximum likelihood phylogenetic analyses resolved a strongly supported clade of *M. melanops*, including the samples from eastern Lake Erie (Fig. 3). There was a relatively deep split (4.9% mean uncorrected p-distance) between one *M. melanops* from the Choctawhatchee River in Florida (JF799448) and the remaining *M. melanops* samples, which exhibit almost no genetic variation despite a broad geographic range spanning Alabama, Louisiana, Wisconsin, Michigan, and New York. This sequence divergence suggests substantial phylogeographic structure and potentially undescribed species-level biodiversity within *M. melanops*.

Fish surveys in the 1800s noted the high abundance of *M. melanops* in western Lake Erie in Ohio and Pennsylvania (Jordan, 1882; Bean, 1893), though the species was thought to be historically absent in the eastern Lake Erie basin of New York (see Carlson et al., 2016; Smith, 1985). Habitat degradation by siltation and channelization from heavy industrialization and agricultural development led to widespread declines of northern *M. melanops* populations in the late 19^th^ and early 20^th^ centuries, including in Lake Erie (White, 1974; Smith, 2002; White and Haag, 1977). By the mid-20^th^ century, *M. melanops* was thought to only be present as “stray” individuals in Ohio tributaries of Lake Erie (White, 1974).

However, in the late 20^th^ and early 21^st^ century, *M. melanops* has increased substantially in abundance in western Lake Erie (Edwards and Staton, 2009; Mandrak and Bouvier, 2014). This increase in abundance was initially hypothesized to be a result of improved sampling effort (Edwards and Staton, 2009) as differences in catchability (i.e., how susceptible a species is to a particular sampling method) typically only varies between species or across life stages, and the total amount of effort (i.e., how much sampling is done) have both been shown to affect the detection rate of fish species during electrofishing surveys (Hangsleben et al., 2013; Lintermans, 2016). While the catchability of *M. melanops* across time is unlikely to have changed, variation in sampling effort could account for the increased observation of this species in eastern Lake Erie drainage. Additionally, in eastern Lake Erie, *M. melanops* has been sporadically observed from the waters of New York starting in the 2000s (see Carlson et al., 2016; Smith, 1985).

To evaluate whether increased detection of *M. melanops* in eastern Lake Erie was the result of an increase in population size or simply the result of increased sampling efforts, we examined electrofishing data from the U.S. Fish and Wildlife Service’s Lower Great Lakes Fish and Wildlife Conservation Office to compare the catch-per-unit-effort rate (i.e., number of fish caught per minute of electrofishing; CPUE) of *M. melanops* in the Buffalo Harbor and Buffalo River since 2012 and 2015, respectively. These data were collected as part of an aquatic invasive species early detection and monitoring program. The focus of those surveys was species identification, presence or absence (detections), and abundance (number caught) of all fish encountered. Therefore, this database can provide some insight of relative abundance changes through time across the areas this program surveys annually. An assessment of the CPUE of *M. melanops* in the Buffalo Harbor and Buffalo River revealed that this species was first detected in this survey in 2021 and was detected again in 2022 (Fig. 4). While this increase in detection rate in the Buffalo River does appear to be associated with an increase in sampling effort, this was not the case for the Buffalo Harbor as over 100 minutes of electrofishing were exerted at this site during 2014 and 2015, and at least 50 minutes of effort was exerted in every year except 2012 and 2017 (Table 4). These results indicate that the recent increase in CPUE of *M. melanops* in the Buffalo Harbor is more likely to be the result of an increase in population density rather than merely the result of increased sampling effort.

**Table 4.** Total U.S. Fish and Wildlife Service Aquatic Invasive Species (AIS) Early Detection and Monitoring electrofishing minutes by year for the Buffalo Harbor, Buffalo River, and the Niagara River.

| **Year** | **Buffalo Harbor** | **Buffalo River** | **Niagara River** |
| --- | --- | --- | --- |
| 2012 | 30 | 0 | 229 |
| 2013 | 50 | 0 | 180 |
| 2014 | 110 | 0 | 180 |
| 2015 | 133 | 30 | 193 |
| 2016 | 70 | 4 | 249 |
| 2017 | 40 | 30 | 310 |
| 2018 | 50 | 20 | 210 |
| 2019 | 80 | 10 | 110 |
| 2021 | 133 | 57 | 261 |
| 2022 | 172 | 101 | 346 |
| Total | 868 | 252 | 2268 |

**Note:** no surveys were conducted in 2020 due to Covid-19 restrictions

**Figure 4.**
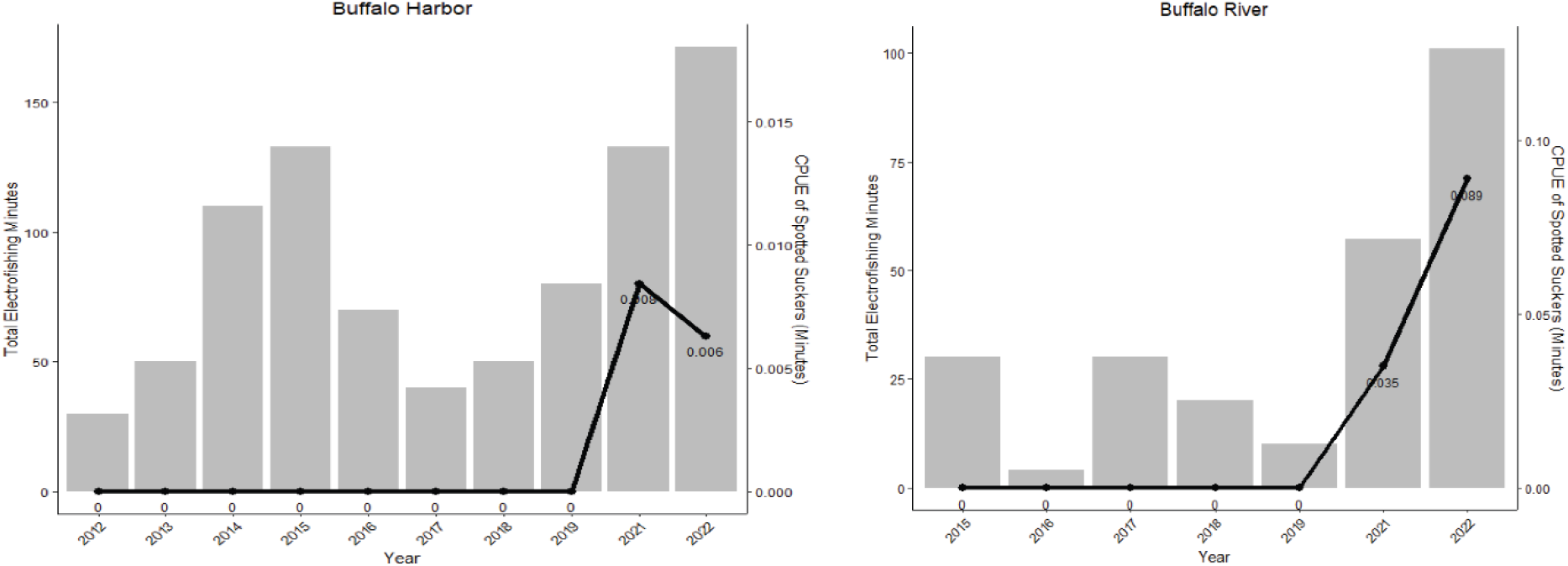
Electrofishing minutes (bars) versus *Minytrema melanops* catch-per-unit-effort (CPUE, lines) by year from the Buffalo Harbor and the Buffalo River. No sampling was conducted in 2020 due to the COVID-19 pandemic.

Like *L. oculatus*, we expect that *M. melanops* will continue its northern and eastern expansion in the Great Lakes basin. *Minytrema melanops* was reported from the Erie Canal in 2007 (NYSM 62256) and from the far eastern shore of Lake Ontario in 2003 (Carlson et al., 2016). Although rare, these sightings suggest that *M. melanops* has already begun dispersing from Lake Erie into Lake Ontario.

### Possible contributing factors to range expansion

The range expansions of *L. oculatus* and the population expansion of *M. melanops* in the eastern basin of Lake Erie may have been facilitated by a combination of climate change and habitat restoration efforts in this region, such as the extensive efforts conducted through the Great Lakes Restoration Initiative (GLRI, www.glri.us). In the Great Lakes, projected increases in air temperature (Hayhoe et al., 2010) are likely to induce warmer water temperatures and longer periods of thermal stratification particularly at shallow depths (Collingsworth et al., 2017). Climate driven range shifts are increasingly commonplace in freshwater fishes, particularly in areas with changing climactic processes that have a higher chance of reporting novel population establishment compared to areas that undergo little change (Fogarty et al., 2017). Rising temperatures in freshwater systems are increasing suitable thermal habitat for warm water fishes by 31% (Mohseni et al., 2003) though there is often a lag between rapid habitat change and responses by impacted species (Gallardo et al., 2016). These range extensions and contractions can lead to varied gains and losses across species within the ecosystem that can ultimately impact community dynamics (Bates et al., 2015). The impacts of climate change on *L. oculatus* and *M. melanops* are not known (see the Fish and Climate Change Database; Krabbenhoft et al. 2020). However, both species are warm water fishes, and therefore northward range expansion of these species is expected to continue as global temperatures rise, similar to expansions observed in other freshwater fishes of Canada and the northern United States (Chu et al. 2005).

In addition to climate change, the Great Lakes have undergone dramatic anthropogenic habitat alteration. Eastern Lake Erie suffered from habitat destruction during the 19^th^ and early 20^th^ centuries, leading to the designation of several areas-of-concern (AOCs; Hartig et al., 2020). At the same time, both *L. oculatus* and *M. melanops* experienced population declines that were attributed to habitat loss. For example, *L. oculatus* appears to have strict habitat preferences and the destruction of these critical habitats has been identified as the primary threat to this species in Ontario, Canada (Glass et al., 2012; Gray et al., 2012). Similarly, habitat alteration has been hypothesized to limit the abundance of lentic prey and the availability of spawning habitat for *M. melanops*, resulting in population declines in the northern portion of its range (Trautman, 1981; White, 1974; White and Haag, 1977). In general, species’ range shifts have been associated with changes in ecological productivity (Ludsin et al., 2001).

Recently, extensive habitat restoration and remediation efforts have been implemented in these waterways (Regier and Hartman, 1973). Habitat restoration efforts in these areas included regulation of pollution (Koonce et al., 1996), shoreline stabilization and naturalization, removal of contaminated substrates, and rebuilding of previously excavated habitat (NOAA, 2022). To this end, the Great Lakes Restoration Initiative has served as an organized attempt at “accelerating efforts to protect and restore the largest system of fresh surface water in the world”, with significant past and ongoing successes. We hypothesize that restoration efforts, along with climate change, have facilitated the range expansion of *M. melanops* and *L. oculatus*, both of which require near-shore habitats for spawning and feeding. Policy makers can use data on native range expansions when considering how to manage lakes with novel disturbances in their ecological networks (Roy et al., 2010). These trends should be considered when managing both native and invasive species, particularly in freshwater systems with recent connectivity that require nuanced estimates of home ranges for habitat conservation plans prioritizing population persistence (Macdonald and Rushton, 2003).

### Conclusions

We reported the first genetically-confirmed records of *L. oculatus* in the eastern basin of Lake Erie in New York, which likely represent an established population. In addition, we reported a population expansion of the *M. melanops* in eastern Lake Erie, supported by genetic evidence. Given that we found no correlation between the increased detection rate of both species and an increase in sampling effort when compared to previous records, we hypothesize that range and population expansion are driven by a combination of climate change and recent habitat restoration. *Lepisosteus oculatus* and *M. melanops* are likely to continue their north and eastward range expansion. Further studies are needed to assess whether these populations are reproducing locally and to characterize population sizes and stability. This study highlights the importance of continued biological monitoring throughout the Great Lakes to better understand changes in aquatic community structure from early detection of non-native species and range shifts of naturally expanding or contracting populations.

## ACKNOWLEDGEMENTS

This work was supported by funds from the Great Lakes Fishery Commission and the University at Buffalo. Fish sampling was conducted under NY DEC scientific collection permit #2442. We thank Doug Carlson, Nick Mandrak, Lisa Holst, John Dettmers, and Jason Robinson for helpful discussions. Corey Krabbenhoft and Levi Gray assisted with field sampling. Daemin Kim and Gregory Watkins-Colwell accessioned YPM ICH.035837 (*L. oculatus*) at the Yale Peabody Museum of Natural History.

## LITERATURE CITED

April, J., Hanner, R.H., Dion-Côté, A.-M., Bernatchez, L., 2013. Glacial cycles as an allopatric speciation pump in north-eastern American freshwater fishes. Mol Ecol 22, 409–422. https://doi.org/10.1111/mec.12116

Bailey, M.R., Smith, G.R., 1981. Origin and Geography of the Fish Fauna of the Laurentian Great Lakes Basin. Can. J. Fish. Aquat. Sci. 38(12), 1539–1561. https://doi.org/10.1139/f81-206

Barbarossa, V., Schmitt, R.J.P., Huijbregts, M.A.J., Zarfl, C., King, H., Schipper, A.M., 2020. Impacts of current and future large dams on the geographic range connectivity of freshwater fish worldwide. Proc. Natl. Acad. Sci. U.S.A. 117, 3648–3655. https://doi.org/10.1073/pnas.1912776117

Bates, A.E., Bird, T.J., Stuart‐Smith, R.D., Wernberg, T., Sunday, J.M., Barrett, N.S., Edgar, G.J., Frusher, S., Hobday, A.J., Pecl, G.T., Smale, D.A., McCarthy, M., 2015. Distinguishing geographical range shifts from artefacts of detectability and sampling effort. Diversity Distrib. 21, 13–22. https://doi.org/10.1111/ddi.12263

Bean, T. H. 1893. The fishes of Pennsylvania: with descriptions of the species and notes on their common names, distribution, habits, reproduction, rate of growth and mode of capture. Harrisburg, PA, E.K. Meyers. https://doi.org/10.5962/bhl.title.33921

Boothroyd, M., Mandrak, N.E., Fox, M., Wilson, C.C., 2016. Environmental DNA (eDNA) detection and habitat occupancy of threatened Spotted Gar (Lepisosteus oculatus): eDNA Detection of Endangered Species. Aquatic Conserv: Mar. Freshw. Ecosyst. 26, 1107–1119. https://doi.org/10.1002/aqc.2617

Broughton, R. E., & Reneau, P. C. 2006. Spatial covariation of mutation and nonsynonymous substitution rates in vertebrate mitochondrial genomes. Molecular Biology and Evolution, 23(8), 1516–1524.

Carlson, D.M., Daniels, R.A., Wright, J., 2016. Atlas of inland fishes of New York. The New York State Education Department and Department of Environmental Conservation.

Chu, C., Mandrak, N. E., & Minns, C. K. 2005. Potential impacts of climate change on the distributions of several common and rare freshwater fishes in Canada. Diversity and Distributions, 11(4), 299–310. https://doi.org/10.1111/j.1366-9516.2005.00153.x

Collingsworth, P.D., Bunnell, D.B., Murray, M.W., Kao, Y.-C., Feiner, Z.S., Claramunt, R.M., Lofgren, B.M., Höök, T.O., Ludsin, S.A., 2017. Climate change as a long-term stressor for the fisheries of the Laurentian Great Lakes of North America. Rev Fish Biol Fisheries 27, 363–391. https://doi.org/10.1007/s11160-017-9480-3

Comte, L., Buisson, L., Daufresne, M., Grenouillet, G., 2013. Climate-induced changes in the distribution of freshwater fish: observed and predicted trends: Climate change and freshwater fish. Freshwater Biology 58, 625–639. https://doi.org/10.1111/fwb.12081

Cudmore-Vokey, B., Crossman, E., 2000. Checklists of the fish fauna of the Laurentian Great Lakes and their connecting channels. Ministère des pêches et des océans.

Dalton, A.S., Margold, M., Stokes, C.R., Tarasov, L., Dyke, A.S., Adams, R.S., Allard, S., Arends, H.E., Atkinson, N., Attig, J.W., Barnett, P.J., Barnett, R.L., Batterson, M., Bernatchez, P., Borns, H.W., Breckenridge, A., Briner, J.P., Brouard, E., Campbell, J.E., Carlson, A.E., Clague, J.J., Curry, B.B., Daigneault, R.-A., Dubé-Loubert, H., Easterbrook, D.J., Franzi, D.A., Friedrich, H.G., Funder, S., Gauthier, M.S., Gowan, A.S., Harris, K.L., Hétu, B., Hooyer, T.S., Jennings, C.E., Johnson, M.D., Kehew, A.E., Kelley, S.E., Kerr, D., King, E.L., Kjeldsen, K.K., Knaeble, A.R., Lajeunesse, P., Lakeman, T.R., Lamothe, M., Larson, P., Lavoie, M., Loope, H.M., Lowell, T.V., Lusardi, B.A., Manz, L., McMartin, I., Nixon, F.C., Occhietti, S., Parkhill, M.A., Piper, D.J.W., Pronk, A.G., Richard, P.J.H., Ridge, J.C., Ross, M., Roy, M., Seaman, A., Shaw, J., Stea, R.R., Teller, J.T., Thompson, W.B., Thorleifson, L.H., Utting, D.J., Veillette, J.J., Ward, B.C., Weddle, T.K., Wright, H.E., 2020. An updated radiocarbon-based ice margin chronology for the last deglaciation of the North American ice sheet complex. Quaternary Science Reviews 234, 106223. https://doi.org/10.1016/j.quascirev.2020.106223

David, S.R., Kik, R.S., Diana, J.S., Rutherford, E.S., Wiley, M.J., 2015. Evidence of countergradient variation in growth of Spotted Gars from core and peripheral populations. Transactions of the American Fisheries Society 144, 837–850. https://doi.org/10.1080/00028487.2015.1040523

David, S.R., Wright, J.J., 2017. Genetic variation and biogeography of the Spotted Gar Lepisosteus oculatus from core and peripheral populations. J. Exp. Zool. (Mol. Dev. Evol.) 328, 596–606. https://doi.org/10.1002/jez.b.22772

DeRoy, E.M., MacIsaac, H.J., 2020. Impacts of invasive species in the Laurentian Great Lakes, in: Crossman, J., Weisener, C. (Eds.), Contaminants of the Great Lakes, The Handbook of Environmental Chemistry. Springer International Publishing, Cham, pp. 135–156. https://doi.org/10.1007/698_2020_593

Ebersole, J.L., Quiñones, R.M., Clements, S., Letcher, B.H., 2020. Managing climate refugia for freshwater fishes under an expanding human footprint. Front Ecol Environ 18, 271–280. https://doi.org/10.1002/fee.2206

Edwards, A.L., Staton, S.K., 2009. Management plan for the Blackstripe Topminnow, Pugnose Minnow, Spotted Sucker and Warmouth in Canada. Species at Risk Act Management Plan Series. Fisheries and Oceans Canada, Ottawa. viii + 43 pp.

Fogarty, H.E., Burrows, M.T., Pecl, G.T., Robinson, L.M., Poloczanska, E.S., 2017. Are fish outside their usual ranges early indicators of climate-driven range shifts? Glob Change Biol 23, 2047–2057. https://doi.org/10.1111/gcb.13635

Gallardo, B., Clavero, M., Sánchez, M.I., Vilà, M., 2016. Global ecological impacts of invasive species in aquatic ecosystems. Glob Change Biol 22, 151–163. https://doi.org/10.1111/gcb.13004

George, S.D., Baldigo, B.P., Collins, S., Clarke, D.B., Winterhalter, D., 2022a. Condition of resident fish communities in the Eighteenmile Creek Area of Concern, New York. Journal of Great Lakes Research 48, 404–411. https://doi.org/10.1016/j.jglr.2020.10.003

George, S.D., Duffy, B.T., Baldigo, B.P., Skaros, D., Smith, A.J., 2022b. Condition of macroinvertebrate communities in the Buffalo River Area of Concern following sediment remediation. Journal of Great Lakes Research 48, 183–194. https://doi.org/10.1016/j.jglr.2021.11.002

Glass, W.R., Corkum, L.D., Mandrak, N.E., 2012. Spring and summer distribution and habitat use by adult threatened Spotted Gar in Rondeau Bay, Ontario, using radiotelemetry. Transactions of the American Fisheries Society 141, 1026–1035. https://doi.org/10.1080/00028487.2012.675904

Glass, W.R., Walter, R.P., Heath, D.D., Mandrak, N.E., Corkum, L.D., 2015. Genetic structure and diversity of Spotted Gar (Lepisosteus oculatus) at its northern range edge: implications for conservation. Conserv Genet 16, 889–899. https://doi.org/10.1007/s10592-015-0708-2

Government of Canada and United States Environmental Protection Agency, 1995. The Great Lakes: An Environmental Atlas and Resource Book. 3rd edition. Toronto, Ontario; Great Lakes National Program Office, U.S. Environmental Protection Agency, Chicago, Illinois. 46 pp.

Gray, S.M., Chapman, L.J., Mandrak, N.E., 2012. Turbidity reduces hatching success in Threatened Spotted Gar (Lepisosteus oculatus). Environ Biol Fish 94, 689–694. https://doi.org/10.1007/s10641-012-9999-z

Great Lakes Water Quality Agreement, 1972. Canada and the United State. Available online: https://www.ijc.org/sites/default/files/C23.pdf. (Accessed Jan 3 2023)

Hangsleben, M.A., Allen, M.S., Gwinn, D.C., 2013. Evaluation of electrofishing catch per unit effort for indexing fish abundance in Florida lakes. Transactions of the American Fisheries Society 142, 247–256. https://doi.org/10.1080/00028487.2012.730106

Hartig, J.H., Krantzberg, G., Alsip, P., 2020. Thirty-five years of restoring Great Lakes Areas of Concern: gradual progress, hopeful future. Journal of Great Lakes Research 46, 429–442. https://doi.org/10.1016/j.jglr.2020.04.004

Hayhoe, K., VanDorn, J., Croley, T., Schlegal, N., Wuebbles, D., 2010. Regional climate change projections for Chicago and the US Great Lakes. Journal of Great Lakes Research 36, 7–21. https://doi.org/10.1016/j.jglr.2010.03.012

Herrington, S.J., Hettiger, K.N., Heist, E.J., Keeney, D.B., 2008. Hybridization between Longnose and Alligator Gars in captivity, with comments on possible gar hybridization in nature. Transactions of the American Fisheries Society 137, 158–164. https://doi.org/10.1577/T07-044.1

Hoang, D.T., Chernomor, O., von Haeseler, A., Minh, B.Q., Vinh, L.S., 2018. UFBoot2: improving the ultrafast bootstrap approximation. Mol Bio and Evo 35, 518–522. https://doi.org/10.1093/molbev/msx281

Hubbs, C., Lagler, K., 2004. Fishes of the Great Lakes Region, revised edition. University of Michigan Press, Ann Arbor, MI. https://doi.org/10.3998/mpub.17658

Inoue, J.G., Miya, M., Tsukamoto, K. and Nishida, M., 2003. Basal actinopterygian relationships: a mitogenomic perspective on the phylogeny of the “ancient fish”. Molecular phylogenetics and evolution, 26(1), pp.110–120.

Jackson, D.A., Peres-Neto, P.R., Olden, J.D., 2001. What controls who is where in freshwater fish communities –the roles of biotic, abiotic, and spatial factors. Can. J. Fish. Aquat. Sci. 58, 157–170. https://doi.org/10.1139/cjfas-58-1-157

Kalyaanamoorthy, S., Minh, B.Q., Wong, T.K.F., von Haeseler, A., Jermiin, L.S., 2017. ModelFinder: fast model selection for accurate phylogenetic estimates. Nat Methods 14, 587–589. https://doi.org/10.1038/nmeth.4285

Katoh, K., Standley, D.M., 2013. MAFFT Multiple Sequence Alignment Software Version 7: improvements in performance and usability. Mol Bio and Evo 30, 772–780. https://doi.org/10.1093/molbev/mst010

Koonce, J.F., Busch, W.-D.N., Czapla, T., 1996. Restoration of Lake Erie: contribution of water quality and natural resource management. Can. J. Fish. Aquat. Sci. 53, 105–112. https://doi.org/10.1139/f96-003

Krabbenhoft, T. J., B. J. E. Myers, J. Wong, C. Chu, R. Tingley Iii, J. A. Falke, T. J. Kwak, C. P. Paukert, A. J. Lynch. 2020. FiCli, Fish and Climate Change Database informs climate adaptation and management for freshwater fishes. Scientific Data. DOI: 10.1038/s41597-020-0465-z

Last, P.R., White, W.T., Gledhill, D.C., Hobday, A.J., Brown, R., Edgar, G.J., Pecl, G., 2011. Long-term shifts in abundance and distribution of a temperate fish fauna: a response to climate change and fishing practices: long-term shifts in a temperate fish fauna. Global Ecology and Biogeography 20, 58–72. https://doi.org/10.1111/j.1466-8238.2010.00575.x

Lintermans, M., 2016. Finding the needle in the haystack: comparing sampling methods for detecting an endangered freshwater fish. Mar. Freshwater Res. 67, 1740. https://doi.org/10.1071/MF14346

Ludsin, S.A., Kershner, M.W., Blocksom, K.A., Knight, R.L., Stein, R.A., 2001. Life after death in Lake Erie: nutrient controls drive fish species richness, rehabilitation. Ecological Applications 11, 731–746. https://doi.org/10.1890/1051-0761(2001)011[0731:LADILE]2.0.CO;2

Lynch, A.J., Cooke, S.J., Deines, A.M., Bower, S.D., Bunnell, D.B., Cowx, I.G., Nguyen, V.M., Nohner, J., Phouthavong, K., Riley, B., Rogers, M.W., Taylor, W.W., Woelmer, W., Youn, S.-J., Beard, T.D., 2016a. The social, economic, and environmental importance of inland fish and fisheries. Environ. Rev. 24, 115–121. https://doi.org/10.1139/er-2015-0064

Lynch, A.J., Myers, B.J.E., Chu, C., Eby, L.A., Falke, J.A., Kovach, R.P., Krabbenhoft, T.J., Kwak, T.J., Lyons, J., Paukert, C.P., Whitney, J.E., 2016b. Climate change effects on North American inland fish populations and assemblages. Fisheries 41, 346–361. https://doi.org/10.1080/03632415.2016.1186016

Lyons, J., Sipiorski, J., 2020. Possible large-scale hybridization and introgression between Longnose Gar (Lepisosteus osseus) and Shortnose Gar (Lepisosteus platostomus) in the Fox River Drainage, Wisconsin. The American Midland Naturalist 183, 105–115. https://doi.org/10.1637/19-043

Macdonald, D.W., Rushton, S., 2003. Modelling space use and dispersal of mammals in real landscapes: a tool for conservation: modelling space use of mammals for conservation planning. Journal of Biogeography 30, 607–620. https://doi.org/10.1046/j.1365-2699.2003.00874.x

Mandrak, N., Bouvier, L., 2014. COSEWIC Status appraisal summary on the Spotted Sucker Minytrema melanops in Canada.

McAllister, K., Drake, D.A.R., Power, M., 2022. Habitat preferences of young-of-year Spotted Gar (Lepisosteus oculatus) in Rondeau Bay, Lake Erie. Can. J. Zool. cjz-2022-0081. https://doi.org/10.1139/cjz-2022-0081

Mcswain, L.E., Gennings, R.M., 1972. Spawning behavior of the Spotted Sucker Minytrema melanops (Rafinesque). Transactions of the American Fisheries Society 101, 738–740. https://doi.org/10.1577/1548-8659(1972)101<738:SBOTSS>2.0.CO;2

Minh, B.Q., Schmidt, H.A., Chernomor, O., Schrempf, D., Woodhams, M.D., von Haeseler, A., Lanfear, R., 2020. IQ-TREE 2: New models and efficient methods for phylogenetic inference in the genomic era. Mol Bio and Evo 37, 1530–1534. https://doi.org/10.1093/molbev/msaa015

Mohseni, O., Stefan, H.G., Eaton, J.G., 2003. Global warming and potential changes in fish habitat in U.S. Streams. Climatic Change 59, 389–409. https://doi.org/10.1023/A:1024847723344

Myers, B.J.E., Lynch, A.J., Bunnell, D.B., Chu, C., Falke, J.A., Kovach, R.P., Krabbenhoft, T.J., Kwak, T.J., Paukert, C.P., 2017. Global synthesis of the documented and projected effects of climate change on inland fishes. Rev Fish Biol Fisheries 27, 339–361. https://doi.org/10.1007/s11160-017-9476-z

Near, T.J., Porterfield, J.C., Page, L.M., 2000. Evolution of cytochrome b and the molecular systematics of Ammocrypta (Percidae: Etheostomatinae). Copeia 2000, 701–711. https://doi.org/10.1643/0045-8511(2000)000[0701:EOCBAT]2.0.CO;2

National Oceanic and Atmospheric Administration, 2022. Current and Past Great Lakes Habitat Restoration Projects, NOAA Fisheries, updated July 18 2022. Available online: https://test-www.fisheries.noaa.gov/national/habitat-conservation/current-and-past-great-lakes-habitat-restoration-projects (accessed Jan 3 2023).

Olden, J.D., Kennard, M.J., Leprieur, F., Tedesco, P.A., Winemiller, K.O., García-Berthou, E., 2010. Conservation biogeography of freshwater fishes: recent progress and future challenges. Diversity and Distributions 16, 496–513. https://doi.org/10.1111/j.1472-4642.2010.00655.x

Ormerod, S.J., Dobson, M., Hildrew, A.G., Townsend, C.R., 2010. Multiple stressors in freshwater ecosystems. Freshwater Biology 55, 1–4. https://doi.org/10.1111/j.1365-2427.2009.02395.x

Page, L.M., Burr, B.M. 2011. Peterson field guide to freshwater fishes of North America north of Mexico, 2nd ed. ed. Houghton Mifflin Harcourt, oston.

Paradis, E., Claude, J., Strimmer, K., 2004. APE: Analyses of phylogenetics and evolution in R language. Bioinformatics 20, 289–290. https://doi.org/10.1093/bioinformatics/btg412

Rahel, F.J., 2002. Homogenization of freshwater faunas. Annu. Rev. Ecol. Syst. 33, 291–315. https://doi.org/10.1146/annurev.ecolsys.33.010802.150429

Regier, H.A., Hartman, W.L., 1973. Lake Erie’s fish community: 150 Years of cultural stresses. Science 180, 1248–1255. https://doi.org/10.1126/science.180.4092.1248

Jordan, D.L., 1882. Report of the Geological Survey of Ohio, Volume 4, Part 1. Columbus, OH. Nevins & Myers

Ricciardi, A., MacIsaac, H.J., 2000. Recent mass invasion of the North American Great Lakes by Ponto–Caspian species. Trends in Ecology & Evolution 15, 62–65. https://doi.org/10.1016/S0169-5347(99)01745-0

Roy, E.D., Martin, J.F., Irwin, E.G., Conroy, J.D., Culver, D.A., 2010. Transient social-ecological stability: the effects of invasive species and ecosystem restoration on nutrient management compromise in Lake Erie. Ecology and Society 15, art20. https://doi.org/10.5751/ES-03184-150120

Scheffer, M., Hosper, S.H., Meijer, M.-L., Moss, B., Jeppesen, E., 1993. Alternative equilibria in shallow lakes. Trends in Ecology & Evolution 8, 275–279. https://doi.org/10.1016/0169-5347(93)90254-M

Scheffer, M., van Nes, E.H., 2007. Shallow lakes theory revisited: various alternative regimes driven by climate, nutrients, depth and lake size. Hydrobiologia 584, 455–466. https://doi.org/10.1007/s10750-007-0616-7

Scheuerell, M.D., Schindler, D.E., 2004. Changes in the spatial distribution of fishes in lakes along a residential development gradient. Ecosystems 7, 98–106. https://doi.org/10.1007/s10021-003-0214-0

Smith, C.L., 1985. The inland fishes of New York State. New York State Department of Environmental Conservation, Albany.

Smith, P.W., 2002. The fishes of Illinois, Published for the Illinois State Natural History Survey by the University of Illinois Press, Urbana, IL.

Snedden, G., Kelso, W., Rutherford, A., 1999. Diel and seasonal patterns of Spotted Gar movement and habitat use in the lower Atchafalaya River Basin, Louisiana. Transactions of the American Fisheries Society 128, 144–154.

Su, G., Logez, M., Xu, J., Tao, S., Villéger, S., Brosse, S., 2021. Human impacts on global freshwater fish biodiversity. Science 371, 835–838. https://doi.org/10.1126/science.abd3369

Suttkus, R., 1963. Order Lepisostei, Fishes of the Western North Atlantic. Yale University Press.

Trautman, M.B., 1981. Fishes of Ohio. Ohio State University Press. https://doi.org/10.2307/j.ctv18bv9k3

Trifinopoulos, J., Nguyen, L.-T., von Haeseler, A., Minh, B.Q., 2016. W-IQ-TREE: a fast online phylogenetic tool for maximum likelihood analysis. Nucleic Acids Res 44, W232–W235. https://doi.org/10.1093/nar/gkw256

Walker, R.H., Kluender, E.R., Inebnit, T.E., Reid Adams, S., 2013. Differences in diet and feeding ecology of similar-sized Spotted (Lepisosteus oculatus) and Shortnose (Lepisosteus platostomus) gars during flooding of a south-eastern US river. Ecol Freshw Fish 22, 617–625. https://doi.org/10.1111/eff.12066

Wheeler, C.R., Gervais, C.R., Johnson, M.S., Vance, S., Rosa, R., Mandelman, J.W., Rummer, J.L., 2020. Anthropogenic stressors influence reproduction and development in elasmobranch fishes. Rev Fish Biol Fisheries 30, 373–386. https://doi.org/10.1007/s11160-020-09604-0

White, D.S., 1974. The biology of Minytrema melanops (Rafinesque), the Spotted Sucker. Ph.D. Dissertation, University of Louisville

White, D.S., Haag, K.H., 1977. Foods and feeding habits of the Spotted Sucker, Minytrema melanops (Rafinesque). American Midland Naturalist 98, 137. https://doi.org/10.2307/2424720

Wright, J.J., David, S.R., Near, T.J., 2012. Gene trees, species trees, and morphology converge on a similar phylogeny of living gars (Actinopterygii: Holostei: Lepisosteidae), an ancient clade of ray-finned fishes. Molecular Phylogenetics and Evolution 63, 848–856. https://doi.org/10.1016/j.ympev.2012.02.033

