## Supplementary Table 1 for "Spotty distributions: Spotted Gar (*Lepisosteus oculatus*) and Spotted Sucker (*Minytrema melanops*) range expansion in eastern Lake Erie"

**Supplementary Table 1** Cytochrome b sequences used for catostomid phylogenetic analysis

| **Species** | **Common name** | **Accession** |
| --- | --- | --- |
| *Carpiodes carpio* | River Carpsucker | AF454867.1 |
| *Carpiodes velifer* | Highfin Carpsucker | JX488762.1 |
| *Carpiodes cyprinus* | Quillback Carpsucker | JX488761.1 |
| *Catostomus commersonii* | White Sucker | KU697932.1 |
| *Catostomus commersonii* | White Sucker | JX488781.1 |
| *Catostomus commersonii* | White Sucker | HQ446762.1 |
| *Deltistes luxatus* | Lost River Sucker | MH603025.1 |
| *Erimyzon succetta* | Lake Chubsucker | AF454878.1 |
| *Erimyzon sucetta* | Lake Chubsucker | KU697910.1 |
| *Erimyzon tenuis* | Sharpfin Chubsucker | AF454877.1 |
| *Erimyzon oblongus* | Creek Chubsucker | AF454876.1 |
| *Hypentelium nigricans* | Northern Hog Sucker | AF454909.1 |
| *Hypentelium roanokense* | Roanoke Hog Sucker | AF454910.1 |
| *Hypentelium etowanum* | Alabama Hog Sucker | AF454908.1 |
| *Ictiobus cyprinellus* | Bigmouth Buffalo | JF799444.1 |
| *Ictiobus niger* | Black Buffalo | JF799446.1 |
| *Ictiobus bubalus* | Smallmouth Buffalo | JF799443.1 |
| *Ictiobus labiosus* | Fleshylip Buffalo | JF799445.1 |
| *Minytrema melanops* | Spotted Sucker | AF454879.1 |
| *Minytrema melanops* | Spotted Sucker | KU697909.1 |
| *Minytrema melanops* | Spotted Sucker | JF799449.1 |
| *Minytrema melanops* | Spotted Sucker | JF799448.1 |
| *Minytrema melanops* | Spotted Sucker | JF799447.1 |
| *Moxostoma duquesnei* | Black Redhorse | AF454895.1 |
| *Moxostoma valenciennesi* | Greater Redhorse | AF454893.1 |
| *Moxostoma robustum* | Robust Redhorse | AF454891.1 |
| *Moxostoma breviceps* | Smallmouth Redhorse | AF454888.1 |
| *Moxostoma carinatum* | River Redhorse | AF454884.1 |
| *Myxocyprinus asiaticus* | Chinese Sucker | AF036176.1 |
| *Thoburnia rhothoeca* | Torrent Sucker | JF799532.1 |
| *Thoburnia rhothoeca* | Torrent Sucker | AF454912.1 |
| *Thoburnia atripinnis* | Blackfin Sucker | AF454911.1 |
| *Xyrauchen texanus* | Razorback Sucker | AF454869.1 |
| *Xyrauchen texanus* | Razorback Sucker | KU697933.1 |
| *Xyrauchen texanus* | Razorback Sucker | JX488825.1 |
