## Supplementary Table 2 for "Spotty distributions: Spotted Gar (*Lepisosteus oculatus*) and Spotted Sucker (*Minytrema melanops*) range expansion in eastern Lake Erie"

**Supplementary Table 2** Cytochrome b sequences used for Lepisosteidae phylogenetic analysis. Asterisks indicate accessions that are likely misidentified.

| **Species** | **Common name** | **Accession** |
| --- | --- | --- |
| *A. spatula* | Alligator gar | NC_008131 |
| *A. spatula* | Alligator gar | JF912043 |
| *A. spatula* | Alligator gar | JF912044 |
| *A. spatula* | Alligator gar | JF912045 |
| *L. oculatus** | Spotted gar | NC_004744 |
| *L. oculatus* | Spotted gar | JF912051 |
| *L. oculatus* | Spotted gar | JF912052 |
| *L. oculatus* | Spotted gar | JF912053 |
| *L. osseus** | Longnose gar | NC_008104 |
| *L. osseus* | Longnose gar | JF912057 |
| *L. osseus* | Longnose gar | JF912058 |
| *L. osseus* | Longnose gar | JF912059 |
| *L. platostomus* | Shortnose gar | JF912054 |
| *L. platostomus* | Shortnose gar | JF912055 |
| *L. platostomus* | Shortnose gar | JF912056 |
| *L. platyrhincus* | Florida gar | JF912048 |
| *L. platyrhincus* | Florida gar | JF912050 |
| *L. platyrhincus* | Florida gar | NC_029715 |
